# Metabolic and Thermal Cues Shape IL-6 Response and Disease Tolerance Mechanisms in Severe Malaria

**DOI:** 10.64898/2026.04.14.718305

**Authors:** Linda Anagu, Samuel Wassmer, Ikenna Anagboso, Jacinta Elo-ilo, Dorothy Ezeagwuna, Alfred Amambua-Ngwa

## Abstract

Severe malaria disproportionately affects children during their earliest Plasmodium falciparum infections, when immunopathology rather than parasite burden often drives clinical deterioration. Because direct investigation of host-parasite interactions during severe disease is ethically impossible, we developed a two-dimensional ex vivo co-culture system that recapitulates key physiological features of malaria pathogenesis. Using PBMCs from malaria-naïve and malaria-exposed adults co-cultured with a freshly adapted P. falciparum isolate, we modelled the combined effects of febrile temperature, pipecolic acid (PA), and lysophosphatidylcholine (LPC) depletion on IL-6 secretion. We also integrated clinical data from children with severe malaria in Anambra State, Nigeria. Across conditions, IL-6 output was not driven by temperature alone but by a metabolically gated interaction: febrile temperature amplified IL-6 only when PA was present, and LPC was not limiting. LPC depletion suppressed IL-6 to near-baseline levels regardless of temperature or PA, indicating that lipid availability constrains inflammatory signalling. Clinical data showed that adverse outcomes clustered with markers of multi-organ dysfunction. Together, these findings support a model in which IL-6 is a context-dependent mediator - participating in inflammatory pathways but not acting as a singular causal driver - and in which metabolic stress, febrile cues, and host tolerance mechanisms jointly shape cytokine production. Ongoing bioinformatics analysis will define the transcriptional responses of both parasite and host cells under these malaria-relevant conditions.

## Introduction

Not everyone infected with *Plasmodium falciparum* will develop severe malaria or die from it. This heterogeneity reflects the early development of malaria tolerance strategies, host mechanisms that regulate overly aggressive immunopathogenic responses during initial infections(1). Severe malaria is therefore most common among children and adults with little or no prior immunity, with the highest risk occurring during the first few infections of life(2). In 2024, malaria caused an estimated 610,000 deaths, predominantly among children in sub-Saharan Africa, including Nigeria (3).

Humans rapidly acquire clinical, non-sterilising immunity to malaria, and immunity increases with age and repeated exposure, resulting in fewer episodes of severe and even uncomplicated malaria (4,5). Naturally acquired clinical immunity involves the regulation of excessive inflammation and is associated with epigenetic reprogramming of innate immune cells, particularly monocytes, during asymptomatic parasitaemia, leading to suppressed cytokine responses (6). In murine malaria models, disease tolerance strategies are deployed early to prioritise host fitness (7). These strategies protect against severe malaria and are acquired quickly following the first infection. Epigenetic reprogramming of monocytes, which prevents their differentiation into inflammatory macrophages, guards against excessive pro-inflammatory responses irrespective of parasite load (6–8).

Innate immune activation during malaria is shaped by parasite-derived and host tissue-damage signals sensed by PBMCs, including hemozoin (9), oxidative stress (10), extracellular vesicles (EVs) (11), and febrile temperatures. EVs from infected erythrocytes act as communication modules that trigger both pro- and anti-inflammatory cytokine responses after being internalised by macrophages (12), potentially contributing to cytokine-storm-like states (11).

Monocytes and macrophages produce pro-inflammatory cytokines such as IL-6 and TNF-α, and both cytokines are associated with poor prognosis in malaria (13,14). Malaria also induces metabolic alterations, including elevated pipecolic acid (PA) in cerebral malaria (15) and depleted lysophosphatidylcholine (LPC) (16). LPC is a signalling molecule that binds Toll-like receptors (17), regulates sexual differentiation in *P. falciparum* (18), and may influence var gene expression through heterochromatin protein 1 (19). PA functions as a signalling molecule in plants (20) and may have analogous roles in host–parasite interactions.

IL-6 is a pleiotropic cytokine involved in innate immunity, with diverse biological activities and complex receptor-mediated signalling pathways (21). Its role in severe malaria remains unclear, but meta-analyses indicate that IL-6 is consistently elevated and may serve as a biomarker of severity (22). However, elevated IL-6 may not be causative (23). IL-6 participates in classic and trans-signalling pathways and influences antigen-specific immune responses and inflammatory reactions (21). Its production during malaria may be triggered by physiological changes: febrile temperatures upregulate *PfSir2b* (24,25) and other virulence-associated pathways (25). Var gene switching among ∼60 var genes is influenced by host immune pressure and environmental cues. In children, severe malaria is often associated with parasites expressing PfEMP1 variants predicted to bind EPCR, disrupting endothelial cytoprotection and impairing disease tolerance.

Children who survive severe malaria often exhibit dampened inflammatory responses, including lower levels of IL-1β, IL-6, and TNF (26). In an ex vivo system mimicking clinical conditions in Malawi, cytokine responses to infected erythrocytes decreased with age. Reduced IL-1β and IL-6 production in children correlated with decreased H3K4me3 at cytokine gene loci (8), highlighting the role of epigenetic reprogramming in survival and immunopathogenesis. These findings underscore the need to understand how pathophysiological factors-temperature, metabolites, lipid availability-shape host survival and parasite virulence.

We hypothesised that altering the host physiological environment would reshape IL-6 responses and influence parasite adaptation. We therefore aimed to explore how these three hallmarks of malaria pathophysiology, febrile temperature, PA augmentation, and LPC depletion, interact to shape IL-6 output and other host and parasite molecular responses. Using a two-dimensional co-culture model with PBMCs from malaria-naïve and malaria-exposed adults co-cultured with a freshly adapted *P. falciparum* isolate, we dissected how environmental perturbations modulate IL-6 secretion. We then linked these mechanistic insights with clinical observations from children with severe malaria in Anambra State, Nigeria.

## METHODS

### Study Site and Design

This mechanistic experimental study was conducted in Anambra State, southeastern Nigeria, where malaria transmission is perennial and peaks during the rainy season from April to October. Locally, malaria is referred to as *Iba*, characterised by fever (*aru okwu* in Igbo). Hospital admissions for malaria peak between July and October, consistent with microscopy-confirmed records from Nnamdi Azikiwe University Teaching Hospital (NAUTH).

We developed an ex vivo two-dimensional co-culture system to mimic key physiological conditions of malaria in both malaria-naïve and malaria-exposed hosts. The primary objective was to quantify IL-6 secretion under malaria-relevant perturbations: reduced blood cell availability, elevated temperature, pipecolic acid (PA) augmentation, and lysophosphatidylcholine (LPC) depletion. A freshly adapted *P. falciparum* field isolate was co-cultured separately with PBMC pools from malaria-naïve adults and malaria-exposed adults. Statistical modelling was used to evaluate the effects of temperature, PA, and LPC on IL-6 output.

### Recruitment and Clinical Samples

#### Children with Severe Malaria

Clinical isolates were obtained from children aged 6 months to <10 years presenting with severe malaria at NAUTH and Okofia Health Centre (OHC). Severe malaria was defined using clinical criteria, including anaemia, impaired consciousness, respiratory distress, multiple convulsions, prostration, shock, abnormal bleeding, pulmonary oedema, and laboratory evidence of jaundice, hypoglycaemia, or renal impairment.

Children were eligible if they were HRP2 mRDT-positive and microscopy-positive for *P. falciparum*. Exclusion criteria included age ≥10 years or lack of parental consent. Recruitment occurred during July-August 2024, the peak admission period. A minimum of five participants was targeted to obtain at least three viable parasite isolates, given an estimated 70% success rate for culture adaptation (27). Ultimately, three isolates were successfully adapted.

#### Adult PBMC Donors

PBMCs were obtained from 12 adult volunteers (22-65 years), with equal numbers of males and females. Six donors were lifelong residents of malaria-endemic Anambra State (malaria-exposed group, PBc), and six were from malaria-free USA (malaria-naïve group, PBn), recruited through Charles River Laboratories.

Eligibility criteria included:

- age ≥18 years
- good general health
- no persistent medical conditions
- negative for *P. falciparum* HRP2
- negative for blood-borne pathogens (Syphilis, CMV, HTLV, West Nile Virus, *Trypanosoma cruzi*, HIV-1/2, Hepatitis B/C)

PBMCs from each group were pooled to generate PBc (immune-experienced) and PBn (immune-naïve) cell populations.

#### Culture Adaptation of *P. falciparum* Isolates

Parasite culture was performed in a certified BL2 malaria culture facility at the Molecular Research Laboratory, NAU Okofia campus, approved by the Nigeria Centre for Disease Control and Prevention (NCDC). The field isolates were cultured in leukocyte-depleted O+ erythrocytes at 4% haematocrit in RPMI-1640–based complete medium containing: 25 µg/mL gentamicin, 0.5% Albumax II, 0.03% glutamine, 0.0026% hypoxanthine, 10% heat-inactivated human serum

Cultures were maintained at 37°C in a tri-gas incubator (5% CO_2_, 5% O_2_, 90% N_2_). Media were replenished every 24-48 hours, and fresh RBCs were added as needed. Parasitaemia and morphology were monitored using Hemacolor® or Giemsa-stained thin smears examined under 100 x oil immersion. Isolates were considered adapted after continuous culture for ≥3 weeks, after which they were cryopreserved in liquid nitrogen.

#### PBMC Isolation and Pooling

PBMCs were isolated using density centrifugation with Histopaque-1077. Whole blood was diluted 1:1 with room-temperature Hanks’ Balanced Salt Solution (HBSS). 35 ml of diluted blood was layered onto 15 ml Histopaque and centrifuged at 400 x g for 30 minutes at 20°C without brake.

Cells at the plasma/Ficoll interface were collected, washed twice in HBSS at 37°C, and centrifuged at 300 x g for 10 minutes. Pellets were combined per donor and resuspended at 2×10^7^ cells/ml in complete RPMI. Aliquots were cryopreserved in freezing medium (50% RPMI, 40% FBS, 10% DMSO) using a controlled-rate freezing container before transfer to liquid nitrogen. Cell counts were performed using trypan blue exclusion and an Improved Neubauer chamber. PBMCs from Nigerian donors (PBc) and U.S. donors (PBn) were pooled separately to generate two biologically distinct immune backgrounds.

#### Co-culture Experiments

Co-culture assays were performed using a freshly adapted *P. falciparum* isolate (Pf Chi) obtained from a child with severe malaria and acute kidney injury. PBMC pools from malaria-exposed donors (PBc) or malaria-naïve donors (PBn) were co-cultured with synchronised ring-stage parasites under malaria-relevant physiological perturbations. A detailed methodology is provided in the [Supplementary file 1](https://1drv.ms/b/c/ba775a116092a823/IQCywj_5GX99QqtNLCr_dFcrAakN--NTl2-SemciYtD_nDM?e=53YDwc).

#### Parasite Synchronisation and Preparation

Parasites were double-synchronised using 5% D-sorbitol in PBS to achieve a synchrony of ± 2 hours. Synchronisation was performed using warm sorbitol solution, and co-culture experiments were initiated when parasites were 7 ± 2 hpi.

#### Experimental Conditions

Co-cultures were established under the following conditions:

- **Temperature:** 37°C (normothermia) or 40°C (febrile range)
- **Pipecolic acid (PA):** 0 or 100µM
- **Lysophosphatidylcholine (LPC):** normal (10% human serum) or depleted (0% serum)
- **Parasite presence:** Pf Chi added vs. no parasite (PBMC-only control)
- **Positive control:** LPS-induced immune activation using DMSO-solubilised LPS

PBMCs and parasites were co-cultured in duplicate wells of a 24-well plate for 8 hours at the assigned temperatures.

#### Post-Culture Processing

After incubation, duplicate wells were pooled into 2 mL RNase/DNase-free tubes. The tubes were centrifuged at 5000 rpm for 5 minutes at room temperature. Then, the supernatants were collected and stored at −20°C for IL-6 ELISA. The pellets were resuspended in 0.8 mL RPMI and layered onto Histopaque in 15 mL tubes. These tubes were centrifuged at 400 g for 30 minutes at 20°C without brake. The PBMC layers and the RBC/platelet pellets were collected separately in tubes. RBC lysis was performed using 0.9 ml RBC lysis buffer on ice for 3 minutes for the PBMC tube. All pellets were washed, centrifuged, and resuspended in 300 µl DNA/RNA Shield for storage at −20°C.

#### IL-6 Quantification by ELISA

IL-6 concentrations in co-culture supernatants were quantified using the AccuBind IL-6 ELISA kit (Monobind Inc., USA). All reagents, standards, and samples were brought to room temperature before use, and the assay was performed strictly according to the manufacturer’s instructions with additional procedural controls to ensure reproducibility. Supernatants were thawed on ice and mixed gently by inversion to avoid protein denaturation. Because preliminary range-finding indicated that IL-6 concentrations fell within the linear range of the assay, samples were analysed without dilution.

Standards were prepared immediately before use by serial reconstitution of the lyophilised calibrator to generate a seven-point standard curve spanning the full dynamic range of the kit. Reference controls supplied by the manufacturer were included on each plate to monitor assay performance. One hundred microlitres of standards, controls, and samples were added to IL-6 antibody-coated microwells in duplicate. Plates were sealed and incubated for the recommended duration to allow antigen binding. After incubation, wells were aspirated and washed thoroughly with the supplied wash buffer to remove unbound material, ensuring that each wash cycle filled and emptied the wells to minimise residual volume.

The anti-IL-6 enzyme conjugate was added to each well, followed by a second incubation period to allow formation of the antibody-antigen-enzyme complex. Plates were washed again to remove excess conjugate. The TMB/H_2_O_2_ substrate solution was then added, and colour development proceeded in the dark at room temperature. Reaction progress was monitored visually to ensure uniform development across wells. The enzymatic reaction was stopped by adding the acidic stop solution, producing a stable yellow endpoint.

Absorbance was measured immediately at 450 nm with a 630 nm reference filter to correct for optical imperfections and plate artefacts. Readings were taken within 15 minutes of adding the stop solution to ensure consistency across plates. Standard curves were generated using a four-parameter logistic (4-PL) model, and IL-6 concentrations in samples were interpolated from the fitted curve. This experiment was done in technical duplicates. Final IL-6 values represent the mean of technical replicates.

#### RNA Extraction, Processing and Sequencing

RNA extraction, quantification, and sequencing were performed by Macrogen Europe.

#### RNA Extraction

Total RNA was extracted using QIAzol® Lysis Reagent and RNeasy® Mini Kit. Up to 1×10^7^ cells were homogenised in 750 µl Trizol, followed by chloroform-mediated phase separation. The aqueous phase was mixed with RLT buffer and ethanol, transferred to RNeasy spin columns under vacuum, washed with RW1 and RPE buffers, dried, and eluted in RNase-free water. RNA concentration was measured using the Quant-IT RiboGreen assay, and integrity was assessed using Agilent TapeStation RNA ScreenTape.

### Library Construction, Preparation and Sequencing

#### Human Samples

Libraries for human RNA-seq were prepared using the TruSeq RNA Access kit (Illumina). Total RNA was first fragmented and converted into first-strand cDNA using SuperScript II, followed by second-strand synthesis incorporating dUTP to maintain strand specificity. The resulting double-stranded cDNA underwent end repair, A-tailing, and adaptor ligation, after which the libraries were enriched by PCR. Hybridisation capture was performed using biotinylated oligonucleotide probes specific to human transcripts, and captured fragments were isolated using streptavidin-coated magnetic beads. A second round of PCR amplification generated the final libraries. Library concentration was quantified using KAPA qPCR assays, and fragment size distribution was assessed using Agilent TapeStation D1000 ScreenTape. Sequencing was carried out on an Illumina NovaSeq X platform using 2×100 bp paired-end chemistry.

#### *Plasmodium* Samples

For *Plasmodium* RNA-seq, libraries were prepared using the SMART-Seq v4 Ultra Low Input RNA kit in combination with the Nextera XT DNA Library Preparation Kit. First-strand cDNA synthesis was performed using template-switching technology to enable full-length transcript capture, followed by purification with SPRI beads. Long-distance PCR amplification generated sufficient cDNA for library construction, and the amplified material was purified using AMPure XP beads. Tagmentation was then performed to fragment and tag the cDNA, after which indexing PCR produced the final libraries. Library concentration was measured by qPCR, and fragment size distribution was evaluated using TapeStation High Sensitivity D5000 ScreenTape. Sequencing was performed on an Illumina NovaSeq platform with 2×100 bp paired-end reads.

#### Bioinformatics Analysis

Bioinformatics analysis was carried out using the nf-core/rnaseq v3.23.0 (28) of the nf-core collection of workflows (29), utilising reproducible software environments from the Bioconda (30) and Biocontainers (31) projects. Quality control (QC), trimming and alignment was done using numerous tools in this pipeline. The pipeline was executed with Nextflow v25.10.4 (32). A gene expression matrix and an extensive QC report were produced in a MultiQC report

A full differential expression and pathway analysis will be included in a subsequent version of this manuscript once de novo assembly and quantification of var genes, differential gene expression analysis, and exploratory data analysis have been finalised.

#### Ethical Approval

Ethical approval (doi:10.6084/m9.figshare.31990104) was obtained from: Nnamdi Azikiwe University Teaching Hospital Ethics Committee (NAUTH/CS/66/VOL.16/VER.3/162/2023/050); extension granted 16 May 2024) and Nnamdi Azikiwe University Human Research Ethics Committee (NAU/HREC/2S/02/08/2023/05).

Written informed consent was obtained from all participants or guardians. Data were de-identified using the Safe Harbour method and will be publicly available upon publication. Participants were informed of their right to withdraw at any time.

### Data Analysis

All analyses were performed in R (version 4.5.0). IL-6 concentrations were log-transformed to stabilise variance and approximate normality. Because PBn cells were pooled and only two technical replicates were obtained per condition, analyses were designed to describe the **pattern and magnitude of effects**, rather than to support subject-level inference.

For the pooled PBn experiment, we modelled log-transformed IL-6 as a function of temperature, pipecolic acid (PA), lysophosphatidylcholine (LPC), and parasite exposure using a linear model of the form:

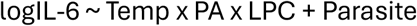

This specification retains the full three-way metabolic interaction (Temp × PA × LPC), while including parasite exposure as an additive term. Higher-order interactions involving the parasite were not estimated because the design used pooled PBn with unbalanced parasite vs no-parasite conditions and no biological replication, making such terms non-identifiable and potentially misleading.

Estimated marginal means (EMMs) for each Temp × PA × LPC combination were obtained using the emmeans package. These EMMs were used to generate: (i) a heatmap of model-adjusted IL-6, (ii) an interaction plot of Temp × PA × LPC, and (iii) a fold-change representation relative to a defined physiological baseline. Because only technical replicates were available, confidence intervals were not plotted for the pooled PBn experiment; figures are presented as model-adjusted means to illustrate the shape and direction of effects. Model diagnostics, including residual plots and normality assessments, are provided in **Supplementary Figure 1 (**[Supplementary file 1](https://1drv.ms/b/c/ba775a116092a823/IQCywj_5GX99QqtNLCr_dFcrAakN--NTl2-SemciYtD_nDM?e=53YDwc)).

Clinical data were summarised using appropriate descriptive statistics and visualised with dot plots overlaid on reference ranges, stratified by outcome categories. No formal multiplicity correction was applied, as the primary aim was mechanistic description rather than hypothesis-testing across multiple endpoints.

More details on the data analysis are in the [Supplementary file 1](https://1drv.ms/b/c/ba775a116092a823/IQCywj_5GX99QqtNLCr_dFcrAakN--NTl2-SemciYtD_nDM?e=53YDwc) (Expanded data analysis).

## RESULTS

### Participants and Clinical Indices

Twelve children with severe malaria were recruited from two health facilities in Anambra State, Nigeria. Ages ranged from 7 months to 8 years, with a slight male predominance. Most cases presented at the tertiary centre (NAUTH), while four presented at Okofia Health Centre (OHC). Prior antimalarial exposure varied: five children had received no treatment before arrival, whereas seven had taken artemisinin-based combination therapy, chloroquine, or herbal preparations. The distribution of cases by site, age, and sex is shown in **Figure 1**.

**Figure 1.**
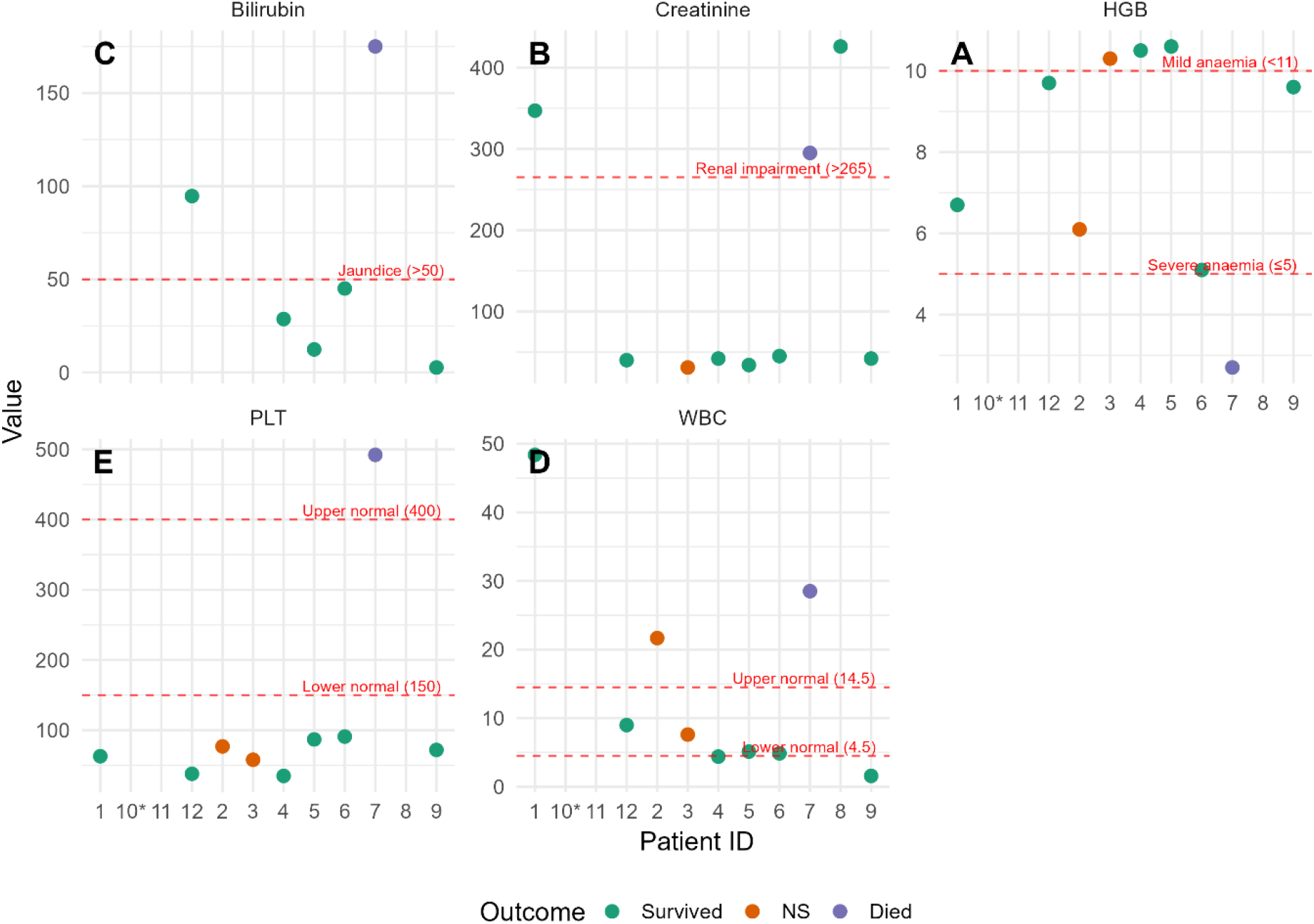
Clinical indices of children with severe malaria, shown as dot-plot panels with clinical reference ranges and outcome-based colour coding. Dot-plot panels showing haemoglobin (HGB), serum creatinine, serum bilirubin, white blood cell count (WBC), and platelet count (PLT) for each child with severe malaria. Horizontal dashed red lines indicate clinically relevant thresholds: severe anaemia (≤5 g/dL), mild anaemia (265 µmol/L), jaundice (>50 µmol/L), normal WBC range (4.5–14.5 ×10^9^/L), and normal platelet range (150–400 ×10^9^/L). Points are colour-coded by clinical outcome (Survived, Neurological Sequelae, Died). This visualisation highlights clustering of multi-organ dysfunction among fatal cases and preserved renal function among children with neurological sequelae.

Eight children survived without neurological sequelae, two survived with neurological sequelae, and two died. Both deaths occurred at NAUTH, representing a 25% case-fatality rate at this site during the two-month recruitment period. One child died within 40 minutes of arrival due to extremely late presentation. Children presenting at OHC generally exhibited less severe symptoms, predominantly severe anaemia, and no deaths occurred at this site.

Laboratory indices at presentation are summarised in **Table 1**. The child who died within three days and had complete laboratory data exhibited profound anaemia (HGB 2.7 g/dL), marked leukocytosis (WBC 28.51×10^9^/L), severe hyperbilirubinaemia (175 µmol/L), hypoglycaemia (42 mg/dL), and extremely elevated creatinine (426 µmol/L), consistent with multi-organ dysfunction. Survivors generally had higher haemoglobin (6–11 g/dL), lower creatinine (31–45 µmol/L), and bilirubin values below 50 µmol/L. Children with neurological sequelae had low haemoglobin but preserved renal function. Platelet counts did not correlate with outcome; thrombocytopenia was common among survivors, whereas the fatal case with complete data had elevated platelets. Although limited by a small sample size, these patterns suggest that severe anaemia, hyperbilirubinaemia, and acute kidney injury at presentation are associated with fatal outcomes, whereas site of care and platelet count are not.

**Table 1:** Patient characteristics, laboratory indices and outcomes.

| ID | Age (months) | Sex | Drug_Hx | Site | Outcome | RGB (mg/dL) | SerumCre (umol/L) | SerumBil (umol/L) | WBC (10e9/L) | Lymph (10e9/L) | Mono (10e9/L) | HGB (g/dL) | HCT (%) | PLT (10e9/L) |
| --- | --- | --- | --- | --- | --- | --- | --- | --- | --- | --- | --- | --- | --- | --- |
| 1 | 7 | F | None | NAUTH | Survived | 94 | 347 | NA | 48.39 | 23.41 | 2.64 | 6.7 | 19.1 | 63 |
| 2 | 36 | F | ACT > 24 h | NAUTH | NS | 99 | NA | NA | 21.67 | 22 | 16.5 | 6.1 | 17 | 77 |
| 3 | 48 | F | None | NAUTH | NS | 98 | 31 | NA | 7.6 | 1.37 | 0.3 | 10.3 | 32.1 | 58 |
| 4 | 48 | M | ACT | OHC | Survived | 102 | 42 | 28.7 | 4.39 | 3.12 | 0.37 | 10.5 | 31.1 | 35 |
| 5 | 60 | M | ACT > 24 h | OHC | Survived | 122 | 34 | 12.4 | 5.12 | 1.82 | 0.76 | 10.6 | 31.5 | 87 |
| 6 | 32 | F | CQ > 24 h | NAUTH | Survived | 160 | 45 | 45.1 | 4.86 | NA | NA | 5.1 | NA | 91 |
| 7 | 19 | M | BAL MIXTU | NAUTH | Died <3 days | 42 | 295 | 175 | 28.51 | 13.9 | 0.62 | 2.7 | 6.9 | 492 |
| 8 | 96 | M | MIXTURES | NAUTH | Survived | 100 | 426 | NA | NA | NA | NA | NA | NA | NA |
| 9 | 24 | M | None | OHC | Survived | 55 | 42 | 2.7 | 1.57 | 0.16 | 0.19 | 9.6 | 29.7 | 72 |
| 10* | 9 | M | None | NAUTH | Died < 40 mins | NA | NA | NA | NA | NA | NA | NA | NA | NA |
| 11 | 11 | M | ACT < 24 h | OHC | Survived | 94 | NA | NA | NA | NA | NA | NA | NA | NA |
| 12 | 63 | F | None | NAUTH | Survived | 66 | 40 | 94.7 | 8.98 | 3.03 | 1.12 | 9.7 | 29.1 | 38 |

Drug\_Hx = drug history, RGB = random blood glucose, SerumCre = serum creatinine, SerumBil = serum bilirubin, Lymph = lymphocytes, Mono = monocytes, PLT = platelets. \*This child was clinically assessed as severely anaemic; grouping and crossmatching for blood transfusion were prioritised. Laboratory results were lost on follow-up as the parent refused to proceed with these tests due to the child's death.

### Bioinformatics Analysis

RNA-seq data have been generated and are undergoing quality control, alignment, and quantification. A full differential expression and pathway analysis will be included in a subsequent version of this manuscript once data analysis has been finalised.

### IL-6 Secretion from Co-culture Supernatants

IL-6 concentrations from PBc–parasite co–cultures were not detectable. IL-6 concentrations from PBn–parasite co–cultures were analysed using a full factorial linear model incorporating temperature, pipecolic acid (PA), and LPC availability. The estimated marginal means (EMMs) for each condition are shown in **Figure 2**, which illustrates the strong dependence of IL-6 output on the interaction between febrile temperature and PA.

**Figure 2.**
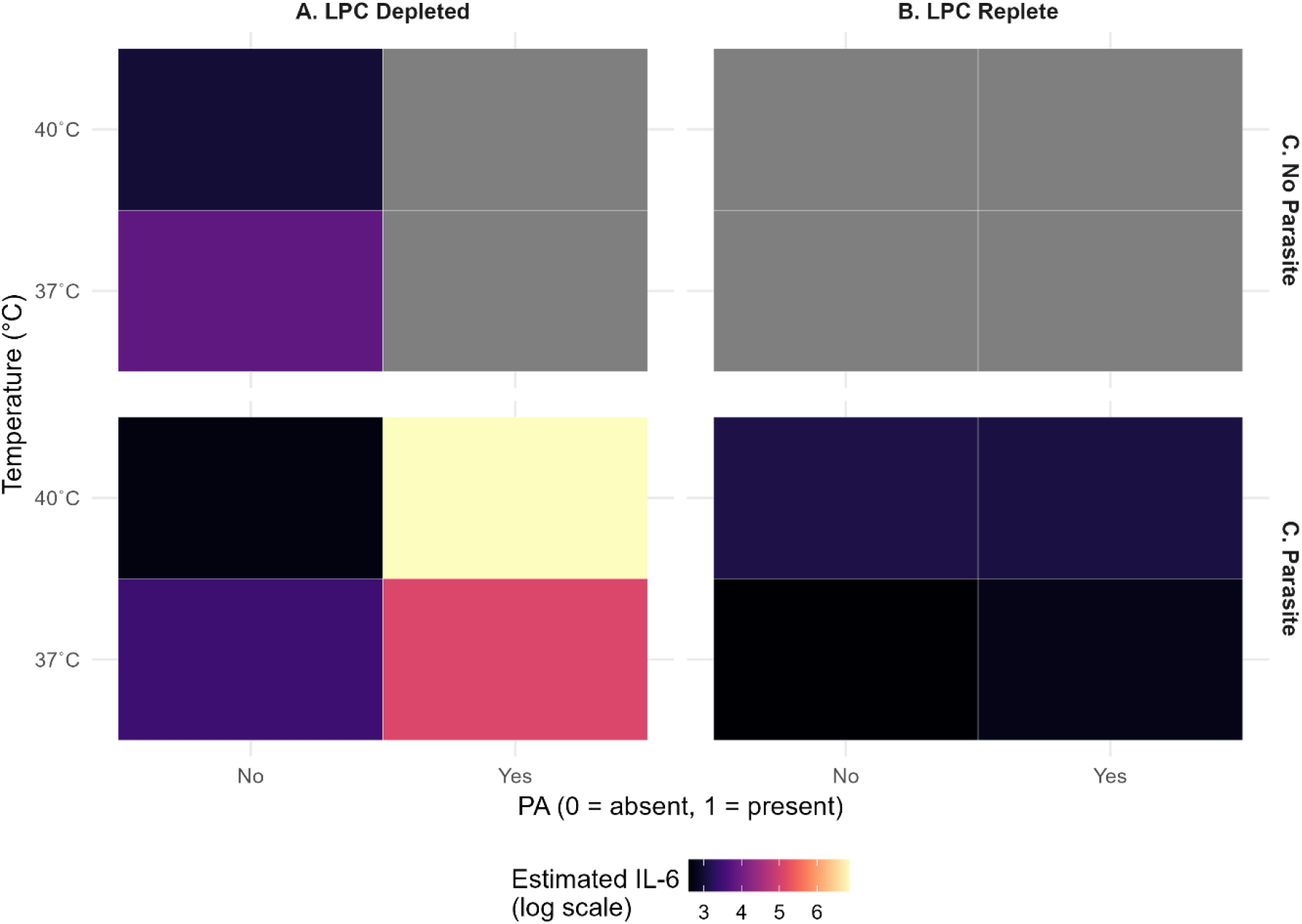
Estimated marginal means of IL-6 across temperature, PA, LPC and parasite conditions. Estimated marginal means (EMMs) of IL‐6 secretion from PBn-parasite co‐cultures under factorial combinations of temperature (37°C, 40°C), pipecolic acid (PA; present/absent), lysophosphatidylcholine (LPC; replete/depleted), and parasite exposure (parasite vs no parasite). Panels A and B show the effects of LPC depletion and repletion, respectively. Panel C shows the effect of parasite exposure independent of metabolic conditions. IL‐6 production is highest under febrile temperature (40°C) when PA is present, and LPC is available. Parasite exposure modestly increases IL‐6 across conditions, but metabolic context remains the dominant determinant of IL‐6 output.

IL-6 secretion increased dramatically at 40°C in the presence of PA, with an approximately 30 to 50-fold rise relative to 37°C without PA. This amplification was not observed when PA was absent, indicating that febrile temperature alone was insufficient to drive a strong IL-6 response. The three-way interaction between temperature, PA, and LPC is visualised in **Figure 3**, which highlights the conditional nature of IL-6 production.

**Figure 3.**
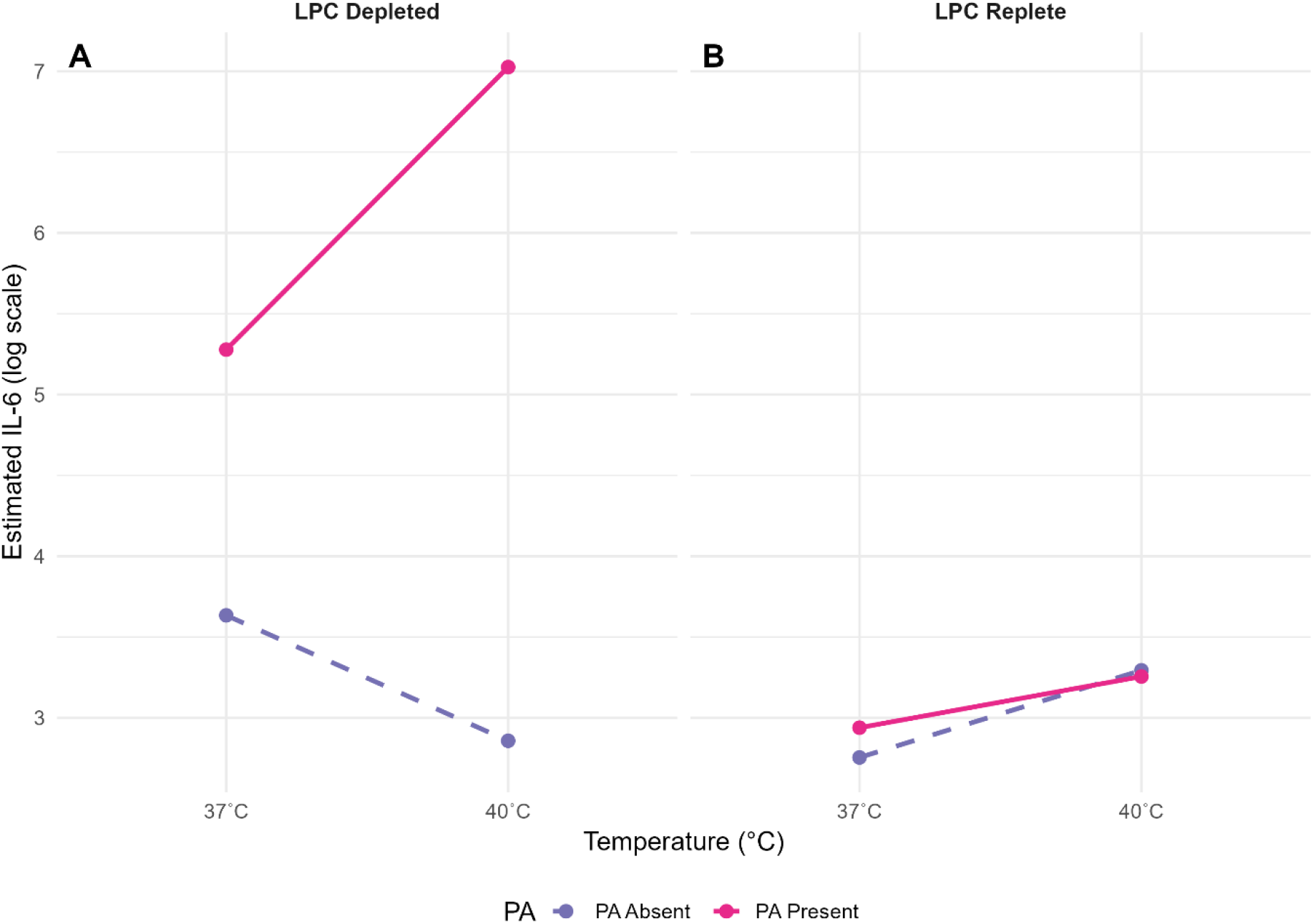
Interaction of Temperature, PA and LPC on IL-6 production. Interaction plot showing estimated marginal means (EMMs) of IL‐6 secretion across factorial combinations of temperature (37°C, 40°C), pipecolic acid (PA; absent/present), and lysophosphatidylcholine (LPC; depleted/replete). Lines represent model‐adjusted means. IL‐6 production increases under febrile temperature (40°C), particularly when PA is present, and LPC is replete. Panel A: LPC‐depleted conditions. Panel B: LPC‐replete conditions.

LPC availability exerted a powerful regulatory effect. When LPC was depleted, IL‐6 concentrations fell to near‐baseline levels across all conditions. Even under the highly stimulatory combination of 40°C and PA, LPC depletion reduced IL‐6 by approximately 40-60‐fold compared with LPC‐replete states, demonstrating that lipid availability is a major constraint on inflammatory signalling in this system. This pattern was evident in both the absolute model‐adjusted means (Figure 2) and in the fold‐change representation (Supplementary Figure 2 in [Supplementary file 1](https://1drv.ms/b/c/ba775a116092a823/IQCywj_5GX99QqtNLCr_dFcrAakN--NTl2-SemciYtD_nDM?e=53YDwc)), where LPC‐depleted conditions remained close to the physiological baseline (37°C, PA absent, LPC replete) regardless of temperature or PA status.

Temperature did not uniformly increase IL-6. Instead, febrile stress acted as an amplifier only when PA was present, and LPC was not limiting. Co-culture with *P. falciparum* modestly elevated IL-6 across conditions, but the dominant drivers of cytokine output were the host-environmental factors rather than parasite presence alone.

Together, these findings reveal a metabolically gated inflammatory response in which IL-6 production depends on the interplay between febrile temperature, infection-associated metabolites, and lipid availability. Rather than behaving as a simple fever-induced cytokine, IL-6 output is conditional on metabolic context, supporting the interpretation that IL-6 is a context-dependent mediator rather than a singular causal driver of severe malaria.

## DISCUSSION

### Metabolic and Thermal Cues Shape IL-6 Output

Severe malaria arises from a complex interplay between parasite replication, host inflammatory responses, and the physiological environment in which these interactions occur. Our findings support a model in which IL-6 secretion is not simply a direct consequence of parasite presence or febrile temperature but is instead shaped by a metabolically gated network of cues that include temperature, infection-associated metabolites, and lipid availability. This aligns with the broader concept of disease tolerance, where host survival depends on limiting tissue damage rather than eliminating parasites (33).

Reduced renal clearance contributes to the elevated pipecolic acid (PA) in children with cerebral malaria due to renal insufficiency (15). However, an enormous amount of PA is generated by the parasite, as measured in culture, rising to 47 µM, and in murine models (34). PA concentrations are equally higher in the brain of mice with neurological abnormalities in an experimental cerebral malaria model, and have been proposed as a putative mediator of neurological dysfunction in this context (15). Current evidence does not demonstrate that PA directly modulates cytokine responses or alters immune activation thresholds in malaria.

Our findings, therefore, extend the existing literature by showing that **PA can directly potentiate IL-6 production under febrile conditions**, revealing a previously unrecognised role for PA as a metabolic amplifier of inflammatory signalling. In this context, PA may act not only as a marker of acute pathology but also as a **functional mediator** capable of shaping early host-parasite interactions when combined with other physiological cues such as temperature and lipid availability.

### LPC Availability as a Constraint on Inflammatory Signalling

Lysophosphatidylcholine (LPC) depletion exerted a profound suppressive effect on IL-6 secretion, reducing cytokine output to near-baseline levels even under highly stimulatory conditions. The plasma of children with malaria contains an assortment of LPCs of varying carbon chain length that are reduced when compared to those of uninfected individuals (35). They play diverse roles in immune signalling, including interactions with Toll-like receptors (17) and regulation of sexual differentiation in *P. falciparum* (18). Its depletion may therefore represent a host- or parasite-driven mechanism to modulate inflammatory tone. The sharp attenuation of IL-6 in LPC-depleted conditions suggests that lipid availability is a critical constraint on cytokine production, consistent with the idea that metabolic resources gate inflammatory responses.

### Reframing IL-6 as a Context-Dependent Mediator

These findings help reconcile the long-standing paradox surrounding IL-6 in malaria. Meta-analyses consistently show elevated IL-6 in severe malaria (22), yet Mendelian randomisation studies indicate that IL-6 signalling is unlikely to be a primary causal driver of severe disease (23), and the evidence of its involvement in malaria severity is currently weak (36). Our data support a blended interpretation: IL-6 participates in the inflammatory cascade and reflects the state of the immune system, but its production is conditional on metabolic and environmental cues. IL-6 is therefore best understood as a context-dependent mediator—both a responder and a signalling molecule within the causal pathway, but not the upstream determinant of severe malaria.

This interpretation is consistent with broader IL-6 biology. IL-6 signals through both classic (membrane-bound IL-6R) and trans-signalling (soluble IL-6R) pathways, each with distinct physiological consequences (21). Selective blockade of IL-6 trans-signalling can suppress inflammation without impairing regenerative functions mediated by classic signalling. The existence of soluble IL-6R and sgp130 as buffering systems further illustrates that IL-6 activity is tightly regulated and highly context-dependent. Our findings extend this paradigm to malaria by demonstrating that IL-6 output is also constrained by metabolic availability, particularly LPC.

### Clinical Data Support a Disease Tolerance Framework

The clinical data from children with severe malaria reinforce the importance of host tolerance mechanisms. Fatal outcomes clustered with markers of multi-organ dysfunction—severe anaemia, hyperbilirubinemia, and acute kidney injury—rather than with any single inflammatory marker. This is consistent with field observations that children who survive severe malaria often exhibit lower circulating levels of pro-inflammatory cytokines (26). Age-stratified analyses from Mali show that monocytes from adults produce lower IL-1β, IL-6, and TNF in response to infected erythrocytes compared to monocytes from children and malaria-naïve adults, and this attenuation is associated with reduced H3K4me3 at cytokine gene loci (8). These findings highlight the role of epigenetic reprogramming in shaping inflammatory trajectories and support the idea that dampened cytokine responses are protective.

### Immune Training and Exposure History Shape IL-6 Potential

Our ex vivo model recapitulates several features of this tolerance landscape. PBc cells (from malaria-exposed adults) produced undetectable IL-6 across conditions, consistent with the regulatory phenotype described in field studies (22,37). In contrast, PBn cells (from malaria-naïve adults) exhibited strong IL-6 responses that were highly sensitive to metabolic and thermal cues. This divergence underscores how prior exposure and immune training shape the inflammatory potential of innate immune cells.

### Parasite Adaptation to Host Cues

The parasite’s ability to sense and respond to host-derived environmental cues is also relevant. Although early studies established PfSir2A and PfSir2B as key regulators of *var* gene silencing (38,39). These did not provide evidence that febrile temperatures upregulate these sirtuins. More recent work demonstrates that heat shock induces a small increase in *PfSir2B* in the trophozoite stage of the parasite (24). Physiologically relevant febrile temperatures ultimately suppress *PfSir2A* and *PfSir2B* expression at the mid-ring stage, leading to altered chromatin structure and increased *var* gene switching (40). Fever, therefore, acts as an epigenetic stressor that can reshape the parasite’s *var* expression landscape.

In children, the severity of *P. falciparum* infection is strongly associated with the expression of *var* genes encoding EPCR-binding PfEMP1 variants, which disrupt endothelial cytoprotection and impair disease tolerance (41). These variants are enriched in severe malaria and are linked to endothelial activation, coagulopathy, and microvascular dysfunction (41–43). Fever-induced modulation of sirtuin activity, combined with metabolic cues such as LPC availability and infection-associated metabolites like PA, may therefore influence the repertoire of PfEMP1 variants expressed during infection. The interplay between host metabolic state, febrile stress, and parasite epigenetic adaptation provides a mechanistic basis for the clinical heterogeneity observed in severe malaria.

### Integrating Mechanistic and Clinical Insights

Overall, our findings support a model in which IL-6 production during malaria is not simply a marker of inflammation but a readout of the host’s metabolic and physiological state. The interaction between temperature, PA, and LPC availability reveals a metabolically gated inflammatory response that aligns with the broader framework of disease tolerance (1). This model provides a mechanistic basis for understanding why some individuals—particularly children with limited prior exposure—mount excessive inflammatory responses that contribute to severe disease, while others maintain a more regulated, tolerance-oriented profile.

## CONCLUSION

This study demonstrates that IL-6 production during malaria is governed by a metabolically gated inflammatory response rather than by parasite presence or fever alone. Metabolic context— particularly the interplay among temperature, infection-associated metabolites, and lipid resources—determines the magnitude of IL-6 output. Immune training and prior exposure shape inflammatory potential, with repeated malaria exposure characterised by minimal IL-6 secretion. Clinical observations from children with severe malaria reinforce this framework: adverse outcomes clustered with multi-organ dysfunction.

By integrating mechanistic, metabolic, and clinical perspectives, this work provides a coherent explanation for why IL-6 is consistently elevated in severe malaria yet unlikely to be a singular causal driver. Instead, IL-6 reflects the physiological state of the host and the metabolic pressures imposed by infection. Understanding how these cues interact offers new insight into malaria pathogenesis and identifies metabolic–immune interfaces as potential targets for future therapeutic or diagnostic strategies.

