## Supplementary material for "Metabolic and Thermal Cues Shape IL-6 Response and Disease Tolerance Mechanisms in Severe Malaria": https://1drv.ms/b/c/ba775a116092a823/IQCywj_5GX99QqtNLCr_dFcrAakN--NTl2-SemciYtD_nDM?e=53YDwc (from another version of this preprint: version 1, 2026-04-16)

**SUPPLEMENTARY FILE 1: Supplementary Methods and Supplementary Figures**

**SUPPLEMENTARY METHODS**

**Co-culture Experiments**

Co-culture experiments were designed to model early host–parasite interactions and to compare inflammatory responses between malaria-naïve (PBn) and malaria-exposed (PBc) adults. All experiments were performed using freshly thawed PBMCs and tightly synchronised *Plasmodium falciparum* ring-stage parasites.

PBMCs were thawed rapidly in a 37°C water bath and immediately transferred into pre-warmed RPMI-1640 supplemented with 10% heat-inactivated human serum. Cells were washed twice by centrifugation at 300 x g for 10 minutes to remove cryoprotectant, then resuspended in the same medium and counted using a haemocytometer or automated counter. Viability was confirmed to be greater than 90% by trypan blue exclusion, and PBMCs were adjusted to a final concentration of $1 \times 10^6$ cells/ml.

Parasites were maintained in continuous culture and synchronised to the ring stage using standard sorbitol treatment. Immediately before co-culture, parasitaemia was quantified on Giemsa-stained thin smears, and cultures were adjusted to 1% parasitaemia and 2% haematocrit. Parasites were washed twice in RPMI-1640 to remove residual metabolites.

Co-cultures were established in 48-well plates by combining PBMCs and parasites at a 1:1 ratio in a final volume of 500 µL per well. Each condition was set up in duplicate. Experimental conditions were generated by varying temperature (37°C or 40°C), the presence or absence of pipecolic acid (PA; added at physiologically relevant concentrations), and LPC availability. For LPC-replete conditions, PBMCs and parasites were cultured in RPMI-1640 supplemented with 10% heat-inactivated human serum. For LPC-depleted conditions, the same RPMI-1640 base medium was used without serum, thereby removing the major exogenous source of LPC. All media and reagents were equilibrated to the target temperature before use.

Plates assigned to the febrile condition were incubated at 40°C in a humidified incubator with 5% $CO_2$, while control plates were maintained at 37°C under identical atmospheric conditions. Temperature stability was monitored throughout the 8-hour incubation. Plates were not disturbed during this period to avoid artefactual cytokine release.

After 8 hours, the co-cultures were processed immediately. Supernatants were collected gently to avoid disturbing the cell pellet, transferred into low-binding tubes, and centrifuged at 400 × g for 5 minutes to remove residual cells and debris. Clarified supernatants were aliquoted into pre-labelled tubes and stored at −80°C until IL-6 quantification. Cell pellets were washed once in ice-cold PBS and processed for RNA extraction or other downstream analyses as required.

All experiments were performed using the same batches of medium and human serum, and parasite synchronisation and PBMC handling were standardised across experimental days. PBMC-only and parasite-only controls were included in each experiment, together with a positive control stimulus, to monitor assay performance and batch variation.

**Expanded data analysis**

**Data structure and preprocessing**

For the pooled PBn experiment, each experimental condition was defined by a factorial combination of:

- **Temperature:** 37 °C (normothermia) or 40 °C (febrile range)
- **PA:** absent vs present (recoded from "No/Yes" to "PA Absent/PA Present")
- **LPC:** replete vs depleted (recoded from "Yes/No" to "LPC Replete/LPC Depleted")
- **Parasite exposure:** parasite vs no parasite

PBn cells were pooled before stimulation, and two technical replicate measurements were acquired for each condition. IL-6 values were log-transformed (natural log) to reduce skew and stabilise variance. Categorical predictors were explicitly recoded as factors with ordered levels to ensure correct model specification and interpretability of EMMs.

**Model specification and rationale**

The final model for IL-6 was:

$$\text{logIL6} \sim \text{Temp} \times \text{PA} \times \text{LPC} + \text{Parasite}$$

This structure was chosen to:

- **Preserve mechanistic focus:** retain the full Temp × PA × LPC interaction to capture how febrile temperature, PA augmentation, and LPC depletion jointly shape IL-6 responses.
- **Avoid over-parameterisation:** exclude the Temp × PA × LPC × Parasite interaction and lower-order parasite interactions, which are not estimable given pooled PBn, unbalanced parasite vs no-parasite conditions, and absence of biological replication.
- **Treat parasite as a contextual modifier:** include parasite exposure as an additive term, allowing estimation of its average effect on IL-6 without implying fully crossed, replicated parasite–metabolic interactions.

Model diagnostics (Supplementary Figure 1) were inspected visually (residual vs fitted plots, Q–Q plots) to confirm that the log-transformed IL-6 data were reasonably compatible with linear model assumptions for descriptive purposes.

**Estimated marginal means and visualisation**

EMMs for each Temp × PA × LPC combination were obtained using emmeans, averaging over parasite exposure. These EMMs underpin:

- **Figure 2:** a heatmap of model-adjusted IL-6 across Temp × PA × LPC, highlighting the dominant contribution of febrile temperature, PA presence, and LPC depletion.
- **Figure 3:** an interaction plot showing EMMs of IL-6 across Temp × PA × LPC, stratified by LPC status (LPC depleted vs replete) and with PA encoded as line colour/linetype. This figure emphasises the shape of the metabolic interaction without confidence intervals, reflecting the technical-replicate design.

For **Supplementary Figure 2**, EMMs were converted to fold-change relative to a physiologically grounded baseline:

- **Baseline:** 37 °C, PA absent, LPC replete (normothermic, PA-low, LPC-replete state).

Fold-change was computed on the log scale as:

$$fold - change = exp(EMM_{condition} - EMM_{baseline})$$

and displayed as a heatmap across Temp × PA × LPC. This representation quantifies how fever, PA augmentation, and LPC depletion shift IL-6 production away from the physiological baseline.

**SUPPLEMENTARY FIGURES**

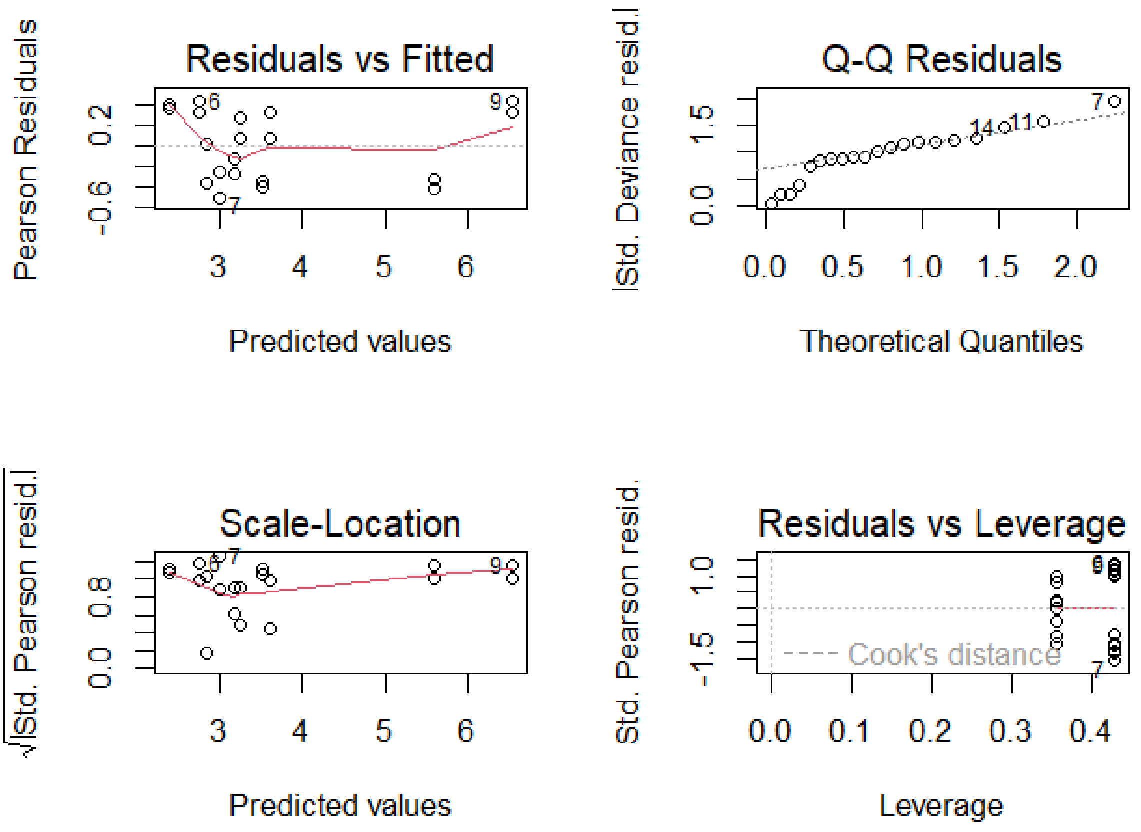


**Supplementary Figure 1. Model diagnostics for IL-6 linear modelling**

Residual plots, Q–Q plots, fitted versus observed values, and leverage assessments demonstrated that model assumptions were met.

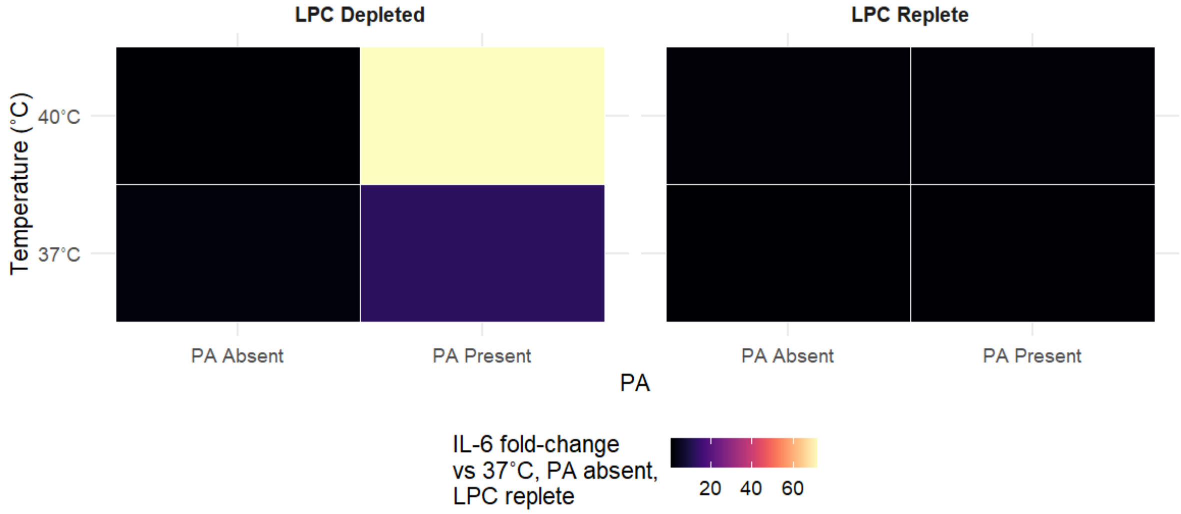


**Supplementary Figure 2. IL-6 fold-change across temperature, PA, and LPC relative to 37˚C, PA absent, LPC replete**

**Supplementary Figure 2.** IL-6 fold-change across temperature, PA and LPC relative to 37˚C, PA absent and LPC replete. IL-6 fold-change across factorial combinations of temperature (37˚C, 40˚C), pipecolic acid (PA; absent/present), and lysophosphatidylcholine (LPC; depleted/replete), relative to the normal physiological baseline (37˚C, PA absent, LPC replete). Fold-change values were derived from model-adjusted means of log-transformed IL-6 using the model logIL6 ~ Temp × PA × LPC + Parasite, averaged over parasite exposure. Fever (40˚C), PA presence, and LPC depletion each increased IL-6 relative to baseline, with the largest fold-changes observed under febrile, PA-present, LPC-depleted conditions.
